# Genetic characterisation of a large antibiotic resistant environmental ST131 *E. coli* plasmid using a long read hybrid assembly approach

**DOI:** 10.1101/2020.06.16.155663

**Authors:** R.S. James, S. Rangama, V. Clark, E.M.H. Wellington

## Abstract

*Escherichia coli* Strain Type 131 are a globally disseminated environmental *E. coli* that has been linked to the capture and spread of plasmid mediated *bla*_CTX-M_ type extended spectrum beta-lactamase (ESBLs). Accurately identifying such resistance genes in their wider genetic context provides a greater understanding of the mechanisms of selection and persistence in the environment. In this study we use a novel DNA extraction and enrichment method in combination with a custom long-read scaffold hybrid-assembly and polishing pipe line to identify the genetic context of the plasmid borne *bla_CTX-M_* gene previously identified in an ST131 environmental *E. coli* isolate. This has allowed us to discern the complete structure of a ~100kb environmental plasmid and further resolve the *bla_CTX-M_* variant to the group 9 *bla_CTX-M-27_* gene. The upstream IS26 insertion element associated with the global capture and dissemination of *bla*_CTX-M-15_ was also identified in proximity to bla_*CTX-M-27*_. Furthermore, the lack of conjugative machinery identified on this plasmid, in combination with a toxin-antitoxin and plasmid partitioning system, indicates a mechanism of vertical transmission to maintain persistence in a population.

## Introduction

The emergence of antibiotic resistance is an ever-expanding global health crisis that is predicted to continue if left unmitigated (Blair et al., 2015, Cassini et al., 2019). Many pathogenic bacteria in both clinical and environmental settings show multi-drug resistant phenotypes which cause untreatable and life-threatening diseases (Obuli, 2016). Conjugative transfer of AMR harbouring plasmids is a major constitute of such antimicrobial resistance (Carattoli, 2013) and furthermore is frequently associated with extra-intestinal pathogenic strains of *E. coli* (Gomi et al., 2017, Manges et al., 2019). Antibiotic resistant strains of extra-intestinal pathogenic *E. coli* (ExPEC) are of particular concern to human health due to the indirect nature of transmission through the environment (Manges et al., 2019, Kaper et al., 2004, Pehrsson et al., 2016). As a result, ExPEC infections are responsible for the majority of the *E. coli* disease burden in locations where sanitation and waste management is either limited (Riley, 2014, Nicolas-Chanoine et al., 2014) or has been disrupted by stochastic environmental events. Given the diverse and global nature of *E. coli* ST131, and the substantial threat posed to human health, understanding the mobility and genetic context of plasmid mediated antibiotic resistance in environmental *E. coli* ST131 is crucial in mitigating the spread of antibiotic resistance in both the developed and the developing world.

Clinical over prescription of broad spectrum third generation cephalosporins (3GCs), such as cefotaxime and ceftazidime, has resulted in wide spread resistance to extended spectrum beta-lactamases (ESBLs). The majority of resistance to 3GCs in pathogenic *E. coli* can be attributed to the beta-lactamase Cefotaxime-Munich (*bla*_CTX-M_) family of resistance genes (Livermore and Hawkey, 2005), with the remainder accounted for by the *bla*_TEM_ and *bla*_OXA_ families. The *bla*_CTX-M_ mediated ESBL resistance is frequently recorded in ExPEC strains of *E. coli* and is particularly associated with the ST131 lineage (Roer et al., 2017, Harris et al., 2018). Group 9 *bla*_CTX-M_ variants are present on mobile genetic elements within *E. coli* ST131, with *bla*_CTX-M-27_ being responsible for upwards of 10% of ESBL resistance in clinical urological *E. coli* infections (Ghosh et al., 2017).

MLST analysis reveals that *E. coli* ST131 is a diverse and globally disseminated pathogenic extra-intestinal *E. coli* (Price et al., 2013) which can exhibit a wide range of plasmid mediated antibiotic resistance profiles. While prolific in nature, one difficulty in understanding the structure and function of such plasmids is the limitation in sequence resolution and assembly given the repetitive nature of low complexity regions and the shared genetic homology between environmental plasmids that co-exist within a host. In this study we aim to develop a method to sequence and resolve the structure and function of a single novel multidrug resistant plasmid hosted by an ST131 *E. coli* (MDR isolate 61) collected from a UK river system. Previous genetic characterisation and antibiotic sensitivity assays has revealed a multi-drug resistance phenotype and the *bla*_CTX-M-27_ genotype, however, the full structure and functions of this plasmid has failed to be resolved using short read data sets. We use a novel plasmid clean up protocol combined with single molecule long read sequencing to resolve the genetic structure of this large environmental plasmid and create a closed sequence for annotation. This has allowed us to determine the genetic context of the emergent *bla*_CTX-M-27_ variant and multidrug resistant gene cassettes and furthermore, provides an insight into the mechanisms of host - plasmid maintenance in a clinically relevant ST131 ExPEC strain found within the UK water system.

## Methods

### DNA Extraction

500 ml of 25ug/ml cefotaxime selective LB broth was inoculated with the *E. coli* isolate and grown overnight in a shaking incubator at 37 °C. DNA was extracted from 500 ml of culture using the Qiagen maxiprep (500) as per manufacturer instructions. DNA was precipitated using room temperature 100% propan −2-ol and spooled onto a closed glass pasture pipette. DNA was dissolved in 1.5 ml 1 x TE buffer at 4°C on a rotating incubator for 1 week. All onwards steps use Eppendorf low bind tubes for sample manipulation.

### Plasmid isolation

Genomic DNA was reduced in the plasmid eluate through degradation with Exonuclease I and Lambda Exonuclease. The reaction of 72 ul consisted of 52 ul of plasmid eluate [2600 ng], 1x lambda exonuclease buffer, 100 units of Exonuclease I and 100 units of lambda exonuclease. Reactions were incubated at 37 °C for fifteen and thirty minutes. Enzymatic degradation was then undertaken at 80 °C for 5 min. 5ul of product was visualised on a 1% agarose gel and the presence of the *bla*_CTX-M_ gene was verified as present in the eluate by endpoint PCR.

### Library preparation

The Oxford nanopore ligation sequencing kit (LSK109) was used to perform the sequencing method. End-repair / dA-tailing and nick repair was undertaken in 60 ul using the ultra II / FFPE repair method. An Oxford nanopore native barcode (BC12) was ligated to the end repaired fragments to increase flow cell efficiency. Barcoded DNA was eluted into 50 μl of H_2_O at 37°C for 15 min. Sequencing adapters were then ligated to barcoded DNA using the NEBNext Quick Ligation module. DNA was eluted into 12ul of ELB (LSK-109) for 15 min at 37 °C.

### MinION sequencing

Sequencing was performed on a MinION MK II. Flow cells were flushed and prepared for sequencing as per manufacturer instructions. MinKNOW V. 2.3 was used to manage the MinION sequencing run. 35.0 μl RBF 25.5 μl Library loading beads and 2.5 μl Nuclease-free water was prepared and mixed before use. Sequencing was left to progress for 2 hours, and then the flow cell was flushed and stored at 4 °C for subsequent use. Real time base calling was disabled and fast5 files were written to an external SSD using RSync and a python shell script. Base calling was undertaken post sequencing using guppy v2.3 and the high accuracy configuration file. Barcodes were demultiplexed and trimmed using guppy_barcoder and qcat.

### Plasmid assembly pipeline

The workflow for this plasmid assembly is outlined in figure 1. Fastq reads were weighted towards a mean quality score of 18 and subsequently down sampled to 500,000 kb using the program Filtlong. Reads shorter than 1.5 kb were not included in this process. De-novo assembly was undertaken using the program Flye with the --plasmid flag enabled. Genome size was estimated at 250 kb for assembly. Contigs were then polished using four rounds of racon and a single round of medaka. Polished contigs were then further improved using existing paired end Illumina data and the program Pilon (figure 1).

**Figure 1.**
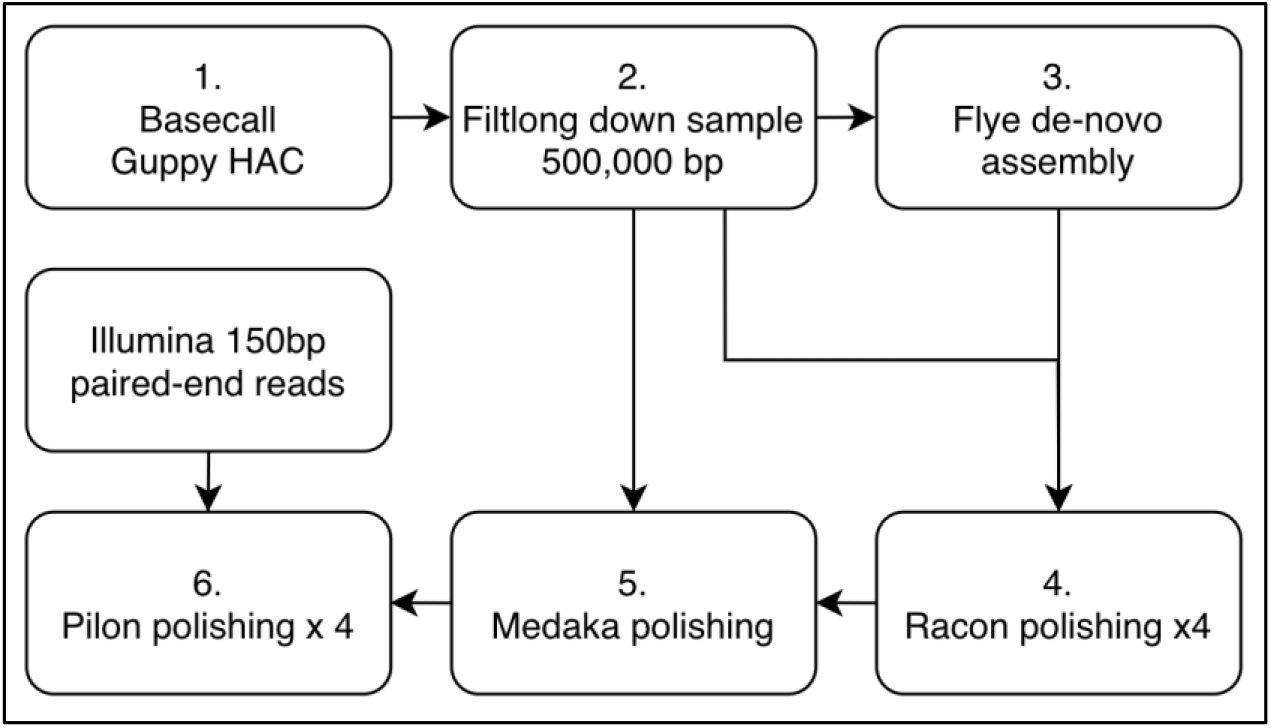
Workflow pipeline for plasmid assembly using both nanopore long-read and Illumina sequence data.

The Ideel pipeline was used to identify and blast query ORFs against the current UniProt protein database (142 GB), then compare identified ORF length with the alignment with the highest identity. The ratio between identified and predicted ORF length was used to give an indication of assembly error with respect to indels that result in falsely truncated or extended ORFs. Plasmid finder v (Carattoli et al., 2014) was used to identify the plasmid incompatibility group at a minimum identity of 95% and a query coverage of 80%. ISfinder was used to identify insertion sequence types in silico with a minimum identity coverage and threshold of 95%.

A custom data base in .gbk format was constructed that contained all annotated sequences on the NCBI data base that corresponded to the query *bla*_CTX-M_ and plasmid or ST131. A total of 237 annotated sequences were used in primary annotation. Open reading frames (ORFs) were identified using Prokka and were annotated using the custom data base as a reference.

The assembled contig was then used to design primers specific to the *bla*_CTX-M_ gene using 100bp flanking regions on the plasmid. Primer design for the *bla*_CTX-M-27_ gene was undertaken in Primer3 (table 1). Long range endpoint PCR was used to amplify the 927bp target sequence. Reactions were undertaken in 25 ul consisting of 1 x Long amp Taq buffer (NEB), 300 μM dNTP mix, 0.4 μM forward and reverse primers, 20 ng BSA, 2.5 U Long Amp Hot Start Taq DNA Polymerase (NEB) and 156 ng DNA template. PCR conditions consisted of 94°C for 2 min, followed by 30 cycles of 94°C for 15 seconds, 56°C for 30 seconds and 65°C for 60 seconds. A final extension at 65°C for 10 min was used. Commercial sanger sequencing (GATC) was then used to sequence the PCR product to identify the *bla*_CTX-M_ variant on the isolated plasmid.

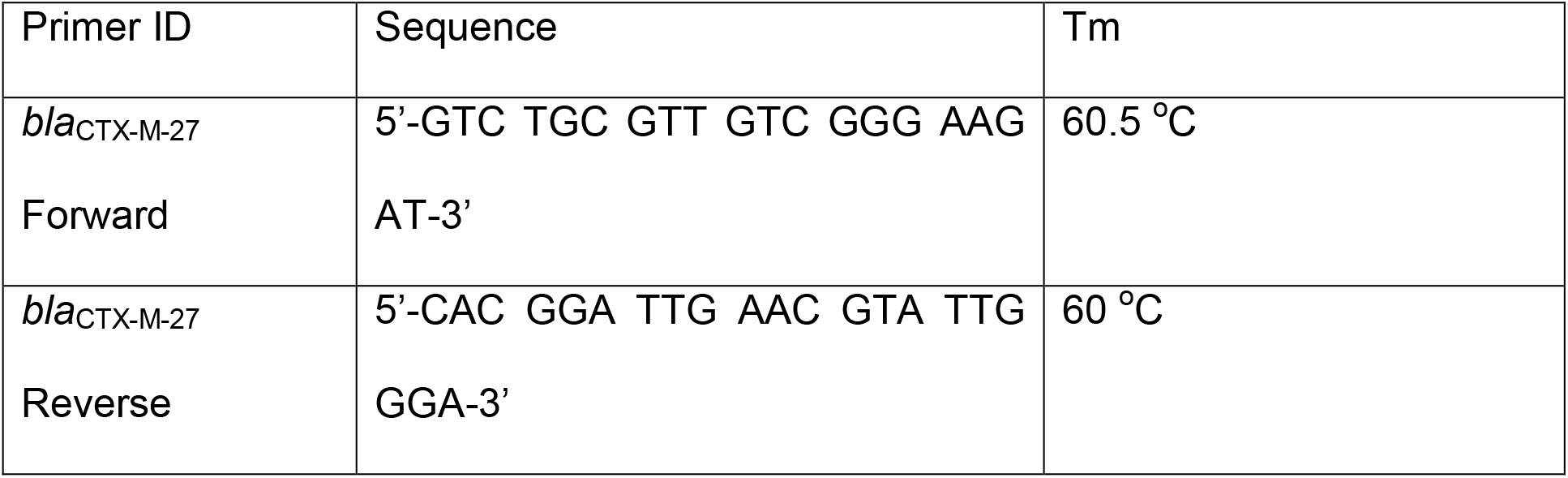

## Results

### DNA extraction and plasmid isolation

Total genomic DNA was recovered in 1.5ml of 1 x TE buffer at a concentration of 156 ng/ul. The majority of DNA recovered was high molecular weight genomic (g) DNA > 10 kb in length (figure 2). The presence of plasmid DNA in covalently closed circular (ccc) open circular (oc) and linear (l) forms was also apparent (figure 2). Linear genomic DNA was substantially degraded following the exonuclease I / Lambda exonuclease digest (figure 2). A total of 1126 ng of DNA was recovered for the sequencing library construction. The presence of *bla*_CTX-M_ was confirmed in the sample using endpoint PCR. The presence of *bla*_CTX-M-27_ was confirmed on the plasmid by Sanger sequencing and referenced against the CARD database at 100% identity and coverage.

**Figure 2.**
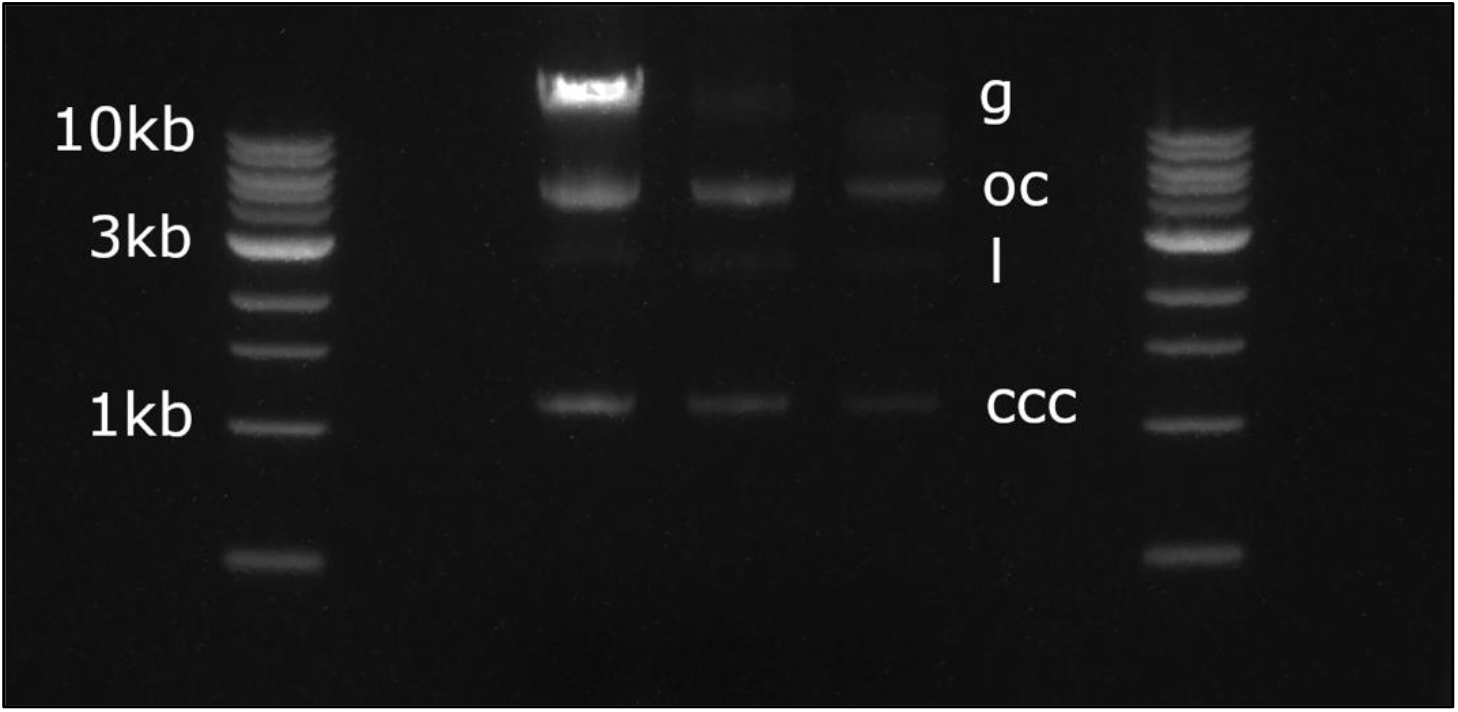
Gel image depicting the HMW DNA extract (Left) and genomic DNA degradation at 15 min (middle) and 30 min (Right). G= genomic liner, oc = open circular, l = liner plasmid, ccc = covalently closed circular plasmid.

### Plasmid sequencing and assembly

Sequencing was undertaken for 2 hours with a total throughput of 1.7 Gb starting with 1611 active pores. Raw read N50 = 17.2 kb with a maximum basecalled read length of 101 kb. A total of one contig was produced from the conclusion of the assembly pipeline with an average coverage of 152 x from long read data and 32 x coverage of Illumina data (5x min cov). Plasmid incompatibility group was identified as IncFII/IncFIA total of 123 ORFs were identified during in the ideel pipeline with <10% having a query-length:reference-length that diverged from 1 (figure 3).

**Figure 3.**
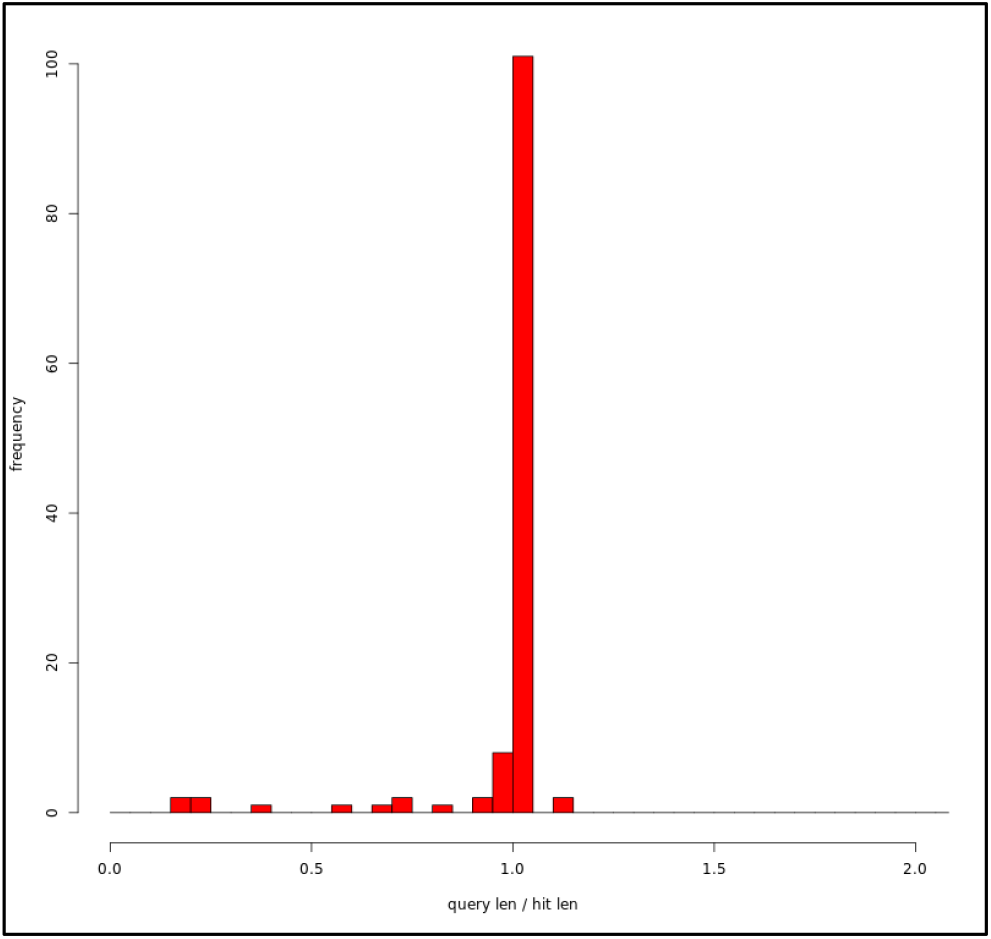
Ratio of predicted ORF length from the assembled and polished plasmid sequence relative to the length of the corresponding hit in the UniProt database. >90% of called ORFs had a query:hit length ratio of 1.

### Plasmid annotation

Automated annotation using Prokka identified a total of 123 ORFs of which 121 were successfully annotated from the reference database (figure 4). A total of 21 ORFs were annotated as hypothetical proteins and were not included in further analysis.

**Figure 4.**
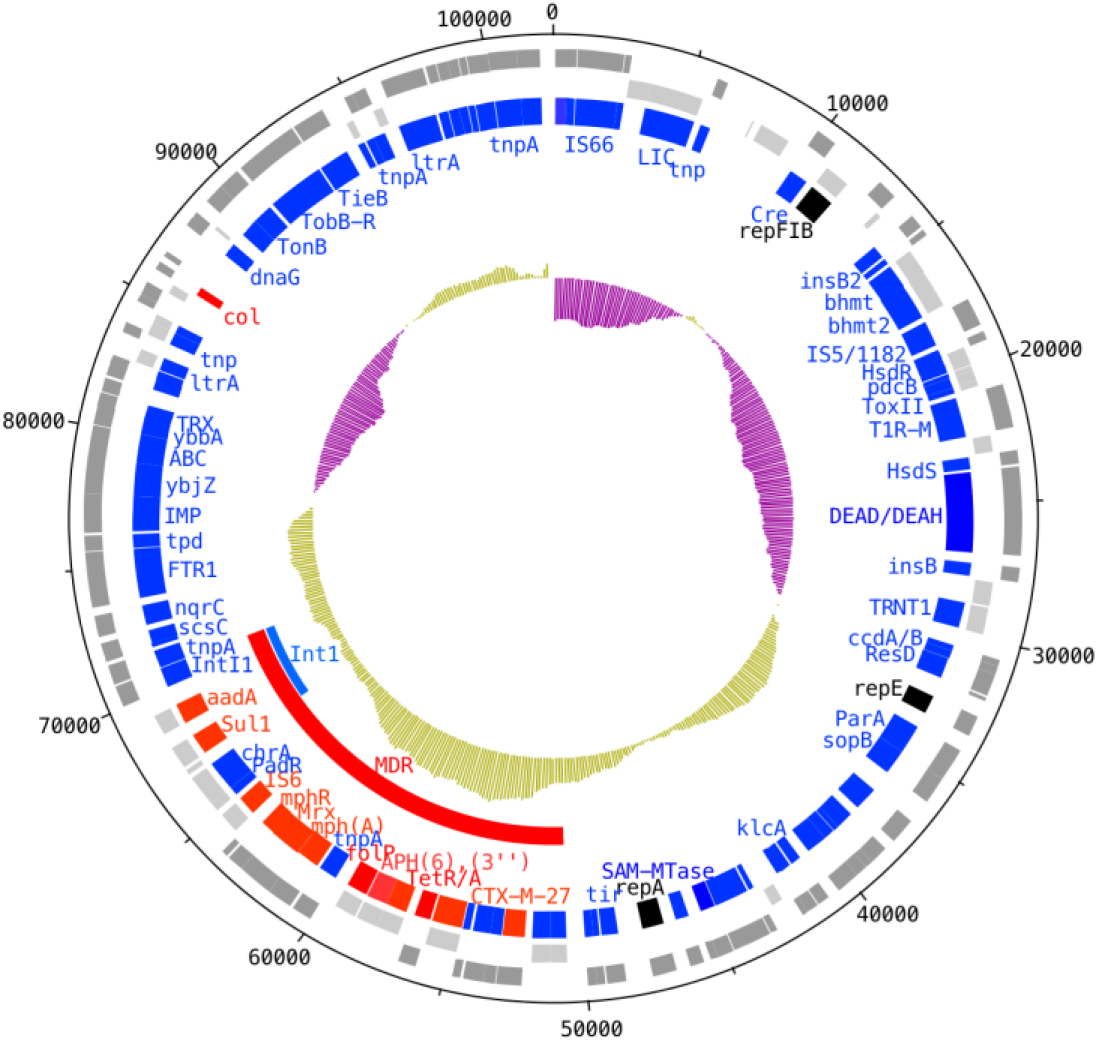
Automated Prokka annotation following plasmid assembly. Inti1 = Class 1 Integron containing aadA and resistance gene, MDR = Multi drug resistance region containing the targeted bla_CTX-M-27_ gene.

Multiple replication regions were identified in this plasmid including RepA, RepFIB and RepE with an overall incompatibility grouping identified by plasmid finder as IncFII / IncFIB at 100% identity and coverage. Nine acquired resistance genes were identified on the plasmid in a multidrug resistant region (figure 3). The *bla*_CTX-M-27_ gene was shown to be present and flanked by tnpA transposase and downstream of an IS26 element at 100% identity and coverage against the ISfinder reference database. The upstream genetic environment was characterised as part of the IS26-ΔISEcp1-blaCTX-M-IS26 type structure (figure 4). A class one integron was also present within the multidrug resistance region upstream of the aminoglycoside resistance gene aadA and sulphonamide resistant gene *sul1*. The MDR region also contained IS6/IS26 and transposase that flank the macrolide resistance genes mphR, Mrx and mph(A). A further resistance gene cassette consisting of the sulphonamide resistance gene *sul2,* two aminoglycoside phosphotransferase genes *APH(6)-Id* and *APH(3”)-Ib* as well as the tetracycline resistance genes *TetA* and associated repressor *TetR*. the IS6 type insertion element ISEc23 was present on the plasmid outside the multidrug resistance region. The plasmid was also shown to possess a type II toxin-antitoxin system as well as genes encoding for the plasmid partitioning proteins ParA and sopB and the ubiquitous ABC transmembrane transporter system. The ccdA/ccdB genes were also present which provide a mechanism for determining the fate of plasmid free daughter cells through replication inhabitation. The *TonB* transport system and associated Colicin gene was also identified on the plasmid as well as the IS66. While three replication regions were identified through in silico analysis, no evidence of conjugative machinery such as the transfer region or pilus construction was found.

## Discussion

The presence of *bla*_CTX-M 27_ was detected on this plasmid by automated annotation of the fully assembled, circularised and annotated sequence. The *bla*_CTX-M 27_ variant was confirmed buy amplification of the target gene and high accuracy Sanger sequencing. This provides strong evidence that the polishing, assembly and annotation method used in this study is reliable given the genetic homogeneity seen within the group 9 CTX-m clade. Furthermore by comparing the length of identified ORFs with their corresponding top hit in the prodigal database it is clear that the proportion of truncated ORFs through sequencing errors is minimal (< 95%) and in in line with the expected proportion of non-functional / psudo genes associated with large environmental plasmids. The reduction in HMW genomic DNA through a novel linear DNA clean up protocol is also apparent and is likely to reduce both genetic contaminations during assembly while increasing sequencing efficiency. This permits a greater resolution of the genetic structure of this plasmid and provides a better understanding of the genetic context of genes of interest such as *bla*_CTX-m 27_, MDR gene cassettes and associated mobile genetic elements.

The role of mobile genetic elements such as transposons and integrons are heavily associated with the horizontal transfer of resistance genes within a bacterial community. The transposable element IS6/IS26 has been linked to the dissemination of both the group 9 and group 1 *bla*_CTX-M_ families, and has been shown to play a decisive role in the capture and spread of *bla*_CTX-M-15_ across the world (Zhao and Hu, 2013). In this plasmid, the *bla*_CTX-M-27_ gene is found within the genetic context of a transposon and directly downstream of an IS26 element and exhibited the conserved genetic environment of IS26-ΔISEcp1-blaCTX-M-like structure associated with the ST131 H30R1 clade of *E. coli* (Matsumura et al., 2015). Mobilization of bla_CTX-M_ genes via an IS located in an upstream environment has been experimentally shown to be a mechanism dissemination at the gene level (Lartigue et al., 2006). Furthermore, the presence of strong promotors associated with insertion sequences upstream are thought to play an important role in AMR gene expression and thus has the potential to strongly influence the selection and mobilisation of this gene in environments under antibiotic selection (Cantón et al., 2012).

Integrons are significant contributors to the problem of antibiotic resistance as they can often be comprised of multi drug resistance gene cassettes that typically confer resistance that originate from different microbial communities. The formation of MDR plasmids can be transferred to a novel host in a single transfer event, resulting in an extremely resistant phenotype. Class I integrons alone are estimated to be present in 40-70% of gram-negative pathogens isolated from both the clinical and environmental settings (Martinez-Freijo et al., 1998) and are often associated with IS6/IS26 in the genetic context of the Group 9 *bla*_CTX-M_ family (Zhao and Hu, 2013).

IncFII plasmids have a relatively narrow host range, are strongly associated with the Enterobacteriaceae and often employ maintenance mechanisms such as tox / anti-tox and ccdA/B systems in order to remain stable within a host population (Bonnin et al., 2012). The presence of a type II toxin-antitoxin system as well as ccdA anti-toxin and and ccdB toxin shows evidence for plasmid stability mechanisms in this environmental strain of *E. coli* by insuring the terminal fate of plasmid free daughter cells that evade vertical transmission. These maintenance mechanisms are thought to explain, in part, the persistence of ESBLs on mobile genetic elements under conditions where antibiotic selection may be low or absent. One way in which a plasmid can further limit the ability of a host to accept novel plasmid is via the acquisition and evolution of different and often multiple replicon types. Replicon types define the replication machinery of the plasmid, which determines the incompatibility (Inc) grouping (Johnson and Nolan, 2009). Plasmids are part of the same incompatibility group if they are unable to stably co-exist in one host, due to use of the same replication machinery or partitioning system (Thomas and Smith, 1987). Many plasmids therefore carry several rep regions decreasing the likelihood another plasmid can invade. In this study, three replication regions were described through in-silico analysis of the plasmid sequence which likely results in exclusion of other large environmental plasmids from invading this host. Group 9 *bla*_CTX-M_ genes are maintained on a number of plasmids of differing incompatibility groupings but are found predominantly on the conjugative IncF and IncK plasmids (Cottell et al., 2011). Both the *bla*_CTX-M-15_ and *bla*_CTX-M-14_ like (group 9) variants are frequently found on multi replicon plasmids with a strong bias towards IncFII plasmids containing FIA/FIB replicons (Naseer and Sundsfjord, 2011).

IncFII plasmids are regarded as being exclusively conjugative in nature and thus are linked to the mobilisation of AMR genes both within and between species of bacteria. However, no evidence of conjugative ability was recorded in this plasmid due to the lack of conjugative machinery detected at the sequence level. It is widely regarded that the *tra* genes associated with conjugative ability are essential for transmission of an IncFIIB plasmid and are responsible for their persistence in a population under selection. The lack of conjugative ability in this plasmid suggests that persistence in maintained solely through toxin / antitoxin systems and inheritance through vertical transmission. This persistence strategy may be advantageous to both the plasmid and the host due to the reduced energy burden associated with conjugation. This has permitted the collection of multi drug resistance genes through site specific insertions and reduced replicative burden due to the presence of multiple replication regions that prevent additional plasmids becoming fixed within a cell line. No evidence was found for the addition of “helper plasmids” that contain separate conjugative machinery which aid in the transfer of larger plasmids between cells. This is the first recorded case of an IncFIIB plasmid in and EXPEC ST131 *E. coli* that maintains persistence in a population without conjugative machinery. The ability to accumulate multiple resistance genes in a single large environmental plasmid hosted by an environmental *E. coli* strain from the ST131 lineage is of clinical relevance due to the potential for pathogenicity and provides a novel in-site into the mechanisms for selective drivers of AMR in the environment.

